# Dual brain cortical calcium imaging reveals social interaction-specific correlated activity in mice

**DOI:** 10.1101/430306

**Authors:** Nicholas J. Michelson, Federico Bolaños, Luis A. Bolaños, Matilde Balbi, Jeffrey M. LeDue, Timothy H. Murphy

## Abstract

We employ cortical mesoscale calcium-imaging to observe brain activity in two head-fixed mice in a staged social touch-like interaction. Using a rail system, mice are brought together to a distance where macrovibrissae of each mouse make contact. Cortical signals were recorded from both mice simultaneously before, during, and after the social contact period. When the mice were together, we observed bouts of mutual whisking and cross-mouse correlated cortical activity in the vibrissae cortex. This correlated activity was specific to individual interactions as the correlations fell in trial-shuffled mouse pairs. Whisk-related global GCAMP6s signals were greater in cagemate pairs during the together period. The effects of social interaction extend outside of regions associated with mutual touch and had global synchronizing effects on cortical activity. We present an open-source platform to investigate the neurobiology of social interaction by including mechanical drawings, protocols, and software necessary for others to extend this work.

## Introduction

The power of social interaction and touch is undisputed across the animal kingdom. In many animals, the presence of a conspecific partner may elicit competitive behavior, reproductive arousal, fear, or other attention-demanding states. Traversal of a social interaction requires each subject to dynamically integrate its internal state and previous experiences with the behavior of its partner and other environmental variables (P. Chen & Hong, 2018). Simultaneously recording neural activity from two individuals engaged in social interaction (Montague et al., 2002) revealed that interacting humans exhibit correlated neural activity (Funane et al., 2011; Liu et al., 2017). Interestingly, this emergent property seems to convey information regarding the context or development of the interaction (Dikker et al., 2017; Jiang et al., 2015; Yang, Zhang, Ni, De Dreu, & Ma, 2020).

Later experiments in mice and bats observed inter-animal neural synchronization at cellular and circuit-level scales, using optical and electrical recording methodologies (Kingsbury et al., 2019; Zhang & Yartsev, 2019). In the mouse prelimbic cortex, single neurons were shown to encode specific self-initiated and partner-initiated competitive social behaviors; and the degree of synchronization between neuronal network activity in each animal was correlated with rank differences in the social dominance hierarchy (Kingsbury et al., 2020). Other studies have shown that prelimbic cortex activity directly modulates social dominance status (Wang et al., 2011; Zhou et al., 2017).

Many circuits throughout the brain shape different aspects of social behavior (Dölen, Darvishzadeh, Huang, & Malenka, 2013; Gunaydin et al., 2014; B. Guo et al., 2019; Rogers-Carter et al., 2018; Sych, Chernysheva, Sumanovski, & Helmchen, n.d.; Tschida et al., 2019; Walsh et al., 2018; Zhou et al., 2017). Given the complex motivational and decision-making states involved with social interaction, it follows that the neural representation of social information may be widespread and distributed throughout the brain similar to other phenomena (Allen et al., 2019, 2017; Pinto et al., 2019; Steinmetz, Zatka-Haas, Carandini, & Harris, 2019). For example, in rodents, information regarding the sex of a conspecific partner is represented in many areas (Ebbesen, Bobrov, Rao, & Brecht, 2019), including prelimbic cortex (Kingsbury et al., 2020), whisker somatosensory cortex (Bobrov, Wolfe, Rao, & Brecht, 2014), and medial amygdala (Li et al., 2017). Moreover, basic sensory signaling is modulated during a social context (Cohen, Rothschild, & Mizrahi, 2011; Lenschow & Brecht, 2015).

Investigation of the macro-scale organization of neural dynamics during social interaction therefore represents an important step forward in understanding the social brain. Widefield cortical calcium imaging provides an opportunity to observe neural activity across the entire dorsal cortex *in vivo* (Clancy, Orsolic, & Mrsic-Flogel, 2019; Gilad & Helmchen, 2020; Musall, Kaufman, Juavinett, Gluf, & Churchland, 2019; Pinto et al., 2019; Vanni, Chan, Balbi, Silasi, & Murphy, 2017; Xiao et al., 2017), but its application to social neuroscience is largely unexplored (MacDowell & Buschman, 2020). In this work, we present a paradigm where multi-subject cortical functional GCaMP imaging is employed during staged interactions between mice. We also provide detailed resources to help investigators set up inexpensive mesoscale cortical GCaMP imaging rigs suitable for dual mouse brain imaging. We find that face to face interactions between mice synchronize cortical activity over wide-scales and this phenomenon is not limited to regions primarily processing whisker/touch dependent signals.

## Methods

### Animals and experimental considerations

All procedures were approved by the University of British Columbia Animal Care Committee and conformed to the Canadian Council on Animal Care and Use guidelines and reported according to the ARRIVE guidelines. Transgenic GCaMP6s tetO-GCaMP6s x CAMK tTA (Wekselblatt, Flister, Piscopo, & Niell, 2016) were obtained from the Jackson Laboratory. All mice used in this study were males >60 days of age and housed in social housing (n=15 mice up to 4 mice/cage from 6 cages) with 12-h light/12-h dark cycles and free access to food and water. We did not employ female mice or male and female mouse pairs because of potential for variation across the estrous cycle that may alter social behavior.

### Surgical procedure

Chronic windows were implanted on male mice that were at least 8 weeks old, as previously described in (Silasi, Xiao, Vanni, Chen, & Murphy, 2016). Fur and skin were removed from the dorsal area of the head, exposing the skull over the entire two dorsal brain hemispheres. After cleaning the skull with a phosphate buffered saline solution, a titanium head-fixing bar was glued to the skull above lambda (Figure 1a) and reinforced with clear dental cement (Metabond). A custom cut coverslip was glued with dental cement on top of the skull (Figure 1a), with the edges of the window reinforced with a thicker mix of dental cement similar to the procedure of (Silasi et al., 2016). Mice recovered for at least seven days prior to imaging or head-fixation.

**Figure 1.**
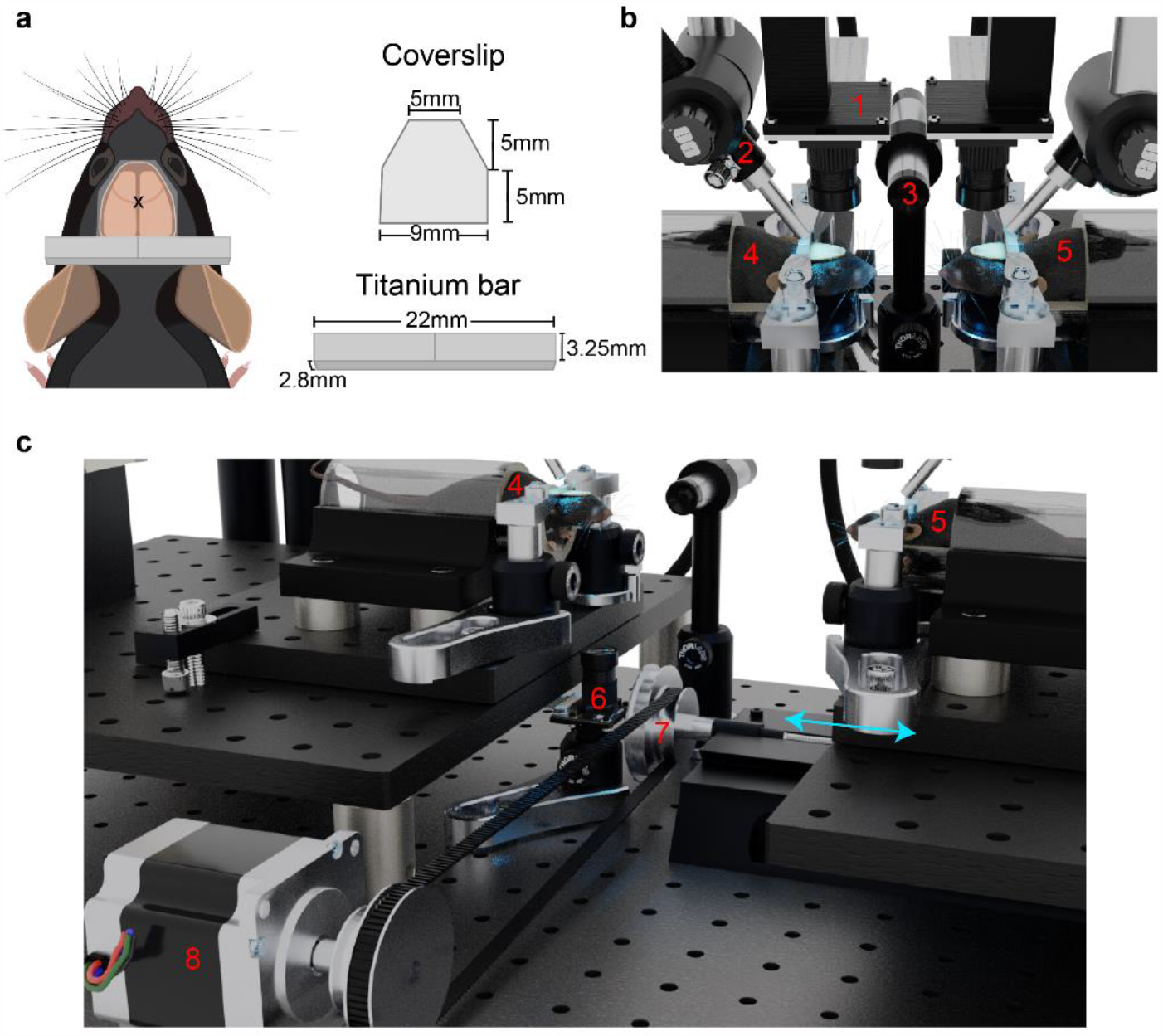
Setup for dual mouse brain imaging system. A) Cartoon depiction of surgical preparation for transcranial mesoscale imaging, with custom cut coverslip and titanium bar for head fixation. B) Close-up view of mouse positioning during the interaction phase of the experiment. C) Larger field of view render of the imaging system. Numbered components are as follows: 1) Raspberry Pi brain imaging camera; 2) GCaMP excitation and hemodynamic reflectance LED light guide; 3) ultrasonic microphone; 4) stationary mouse; 5) moving mouse; 6) Raspberry Pi infrared behavior camera; 7) stage translation knob; 8) stepper motor with belt controlling stage translation. Blue arrows indicate direction of motion of the translatable rail.

### Social Dominance Measurements

Social rank was estimated using the tube-test assay (Fan et al., 2019). Briefly, mice were introduced to either end of a narrow plexiglass tube (32cm long, 2.5cm inner diameter). Upon meeting in the middle, mice compete by pushing each other to get to the opposite side. The mouse which pushes the other back out of the tube is deemed the winner. All combinations of mice within a cage were tested in a round robin format to determine the linear dominance hierarchy. Tube test tournaments were repeated weekly to assess stability of the hierarchy.

### Social Interaction Experiments

Two Raspberry Pi imaging rigs were set up facing each other, and initially separated by 14 cm. A parts list and assembly instructions for the Raspberry Pi widefield imaging rig are included in the supplementary information. One imaging rig was placed atop a translatable rail (Sherline 5411 XY Milling Machine Base), which was driven by a stepper motor to bring the mouse (hereafter referred to as the moving mouse) into the proximity of the other mouse (stationary mouse, 6-12 mm inter-snout distance) (see Table 1 and supplemental build guide for details). Thus, we imaged dorsal cortical activity from two head-restrained mice simultaneously, while varying the distance between snouts (Figure 1b-c, Supplemental Video 1). Mice were habituated to the system for at least one week prior to conducting experiments by head-fixing the animals each day and performing all procedures (e.g. translation, imaging) without the presence of the other mouse.

**Table 1:**
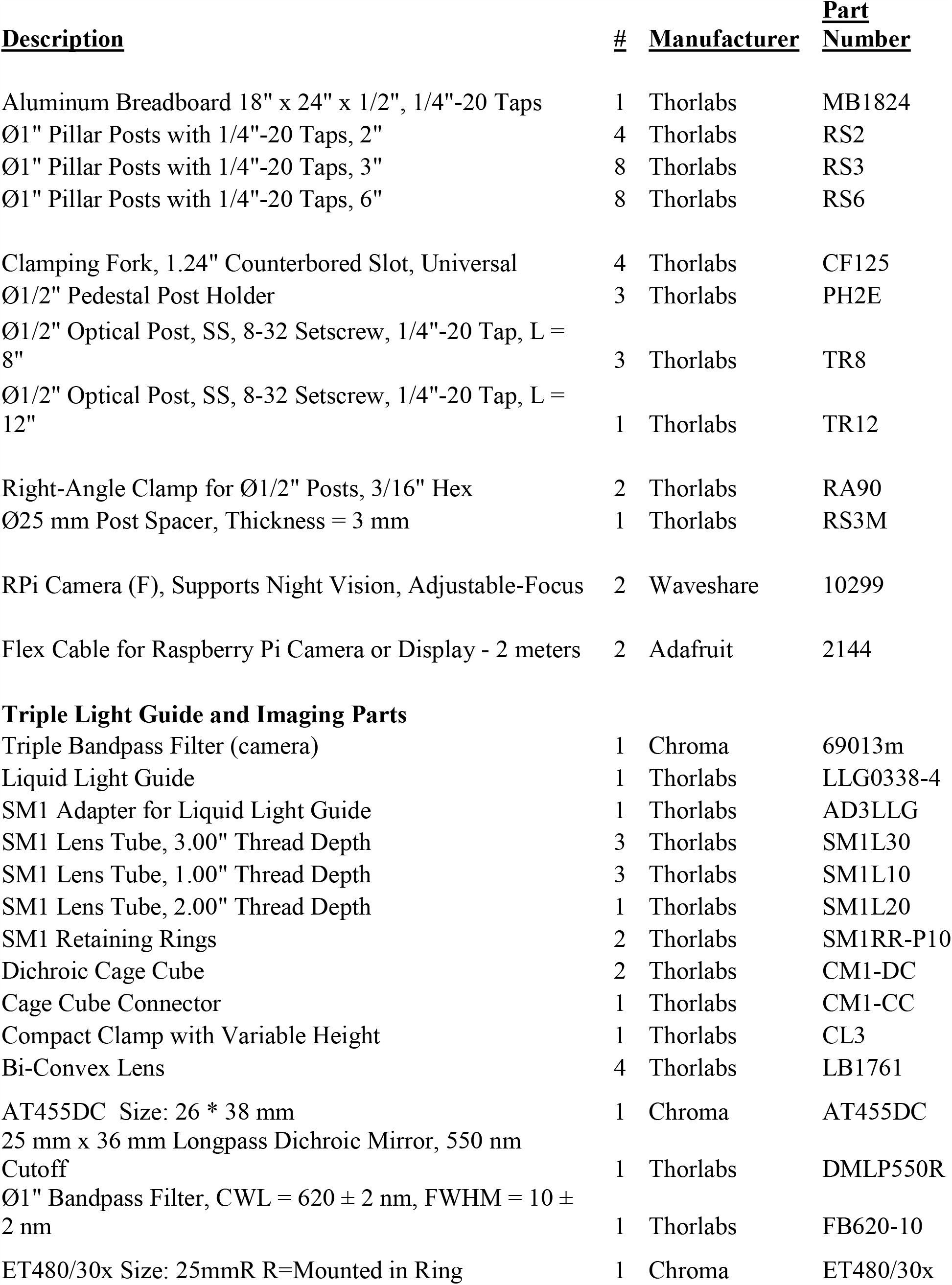
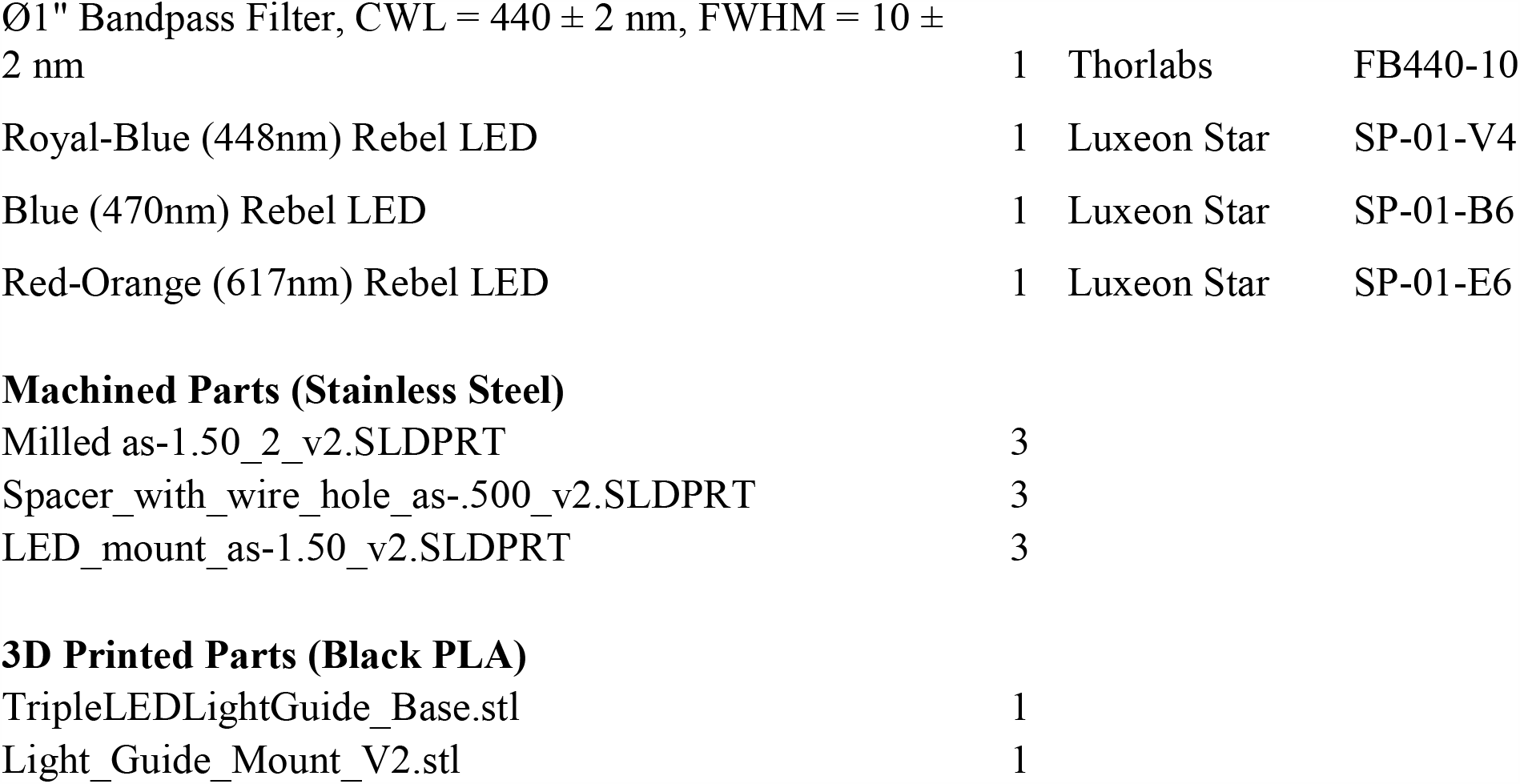
Parts List for Social Interaction System.

The entire imaging system was housed inside a box lined with acoustic foam, thereby reducing ambient light and noise. Throughout the experiment, audio recordings were obtained at 200 kHz using an ultrasonic microphone (Dodotronic, Ultramic UM200K) positioned within the recording chamber approximately 5cm from each mouse’s snout. Audio recordings were analyzed for ultrasonic vocalizations using the MATLAB toolbox DeepSqueak (Coffey, Marx, & Neumaier, 2019).

### Behavior Imaging

The experimental setup was illuminated with an infrared (850 nm) light emitting diode (LED) and behaviors were monitored using an infrared Raspberry Pi camera (OmniVision, OV5647 CMOS sensor). Behavior videos were captured at a framerate of 90 frames per second (fps) with a resolution of 320×180 pixels. The camera was positioned such that the stationary mouse was always included in the field of view and both mice were clearly visible when they were together (Figure 2a).

**Figure 2.**
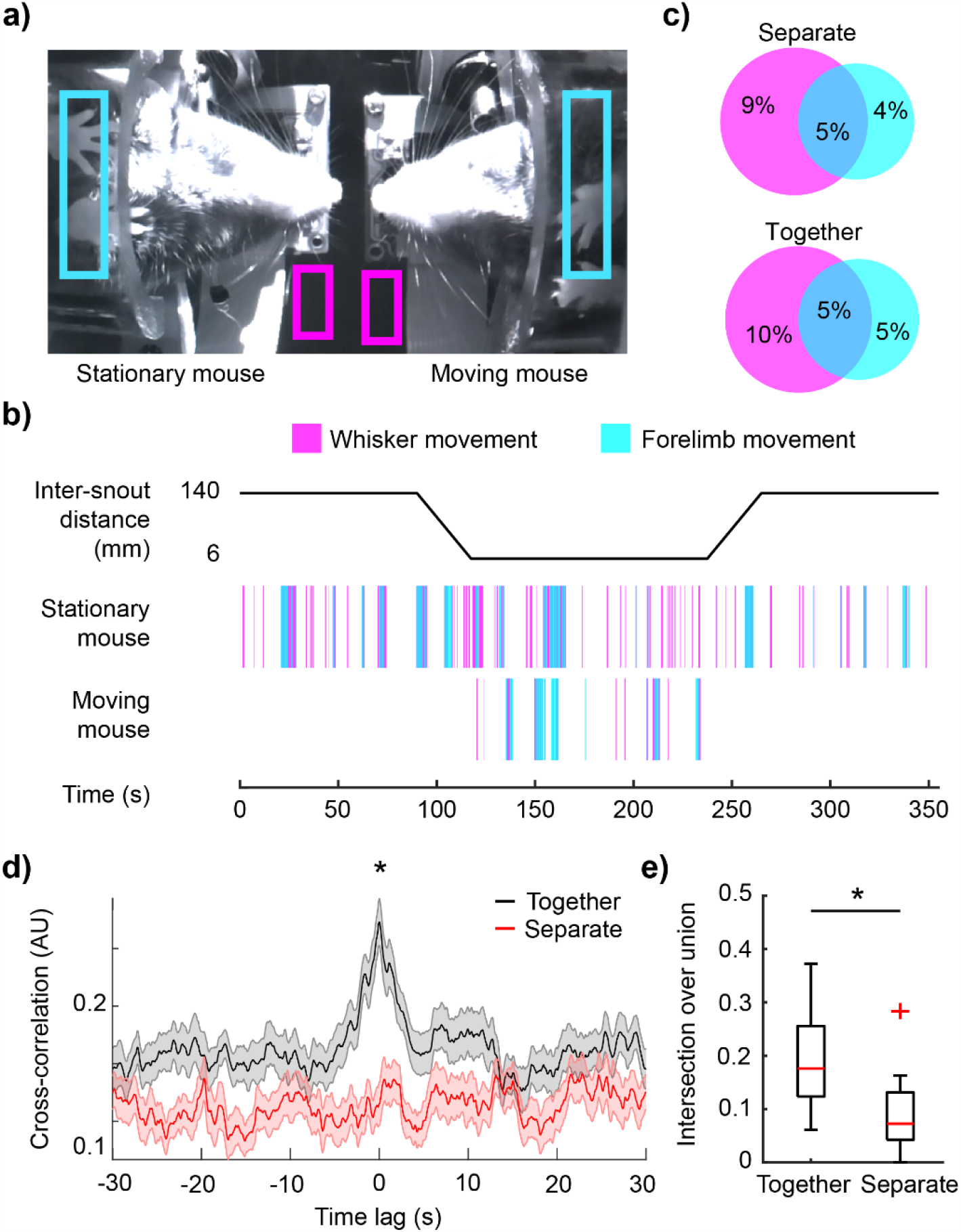
Mice coordinate behavior during interaction. A) Example image of both mice during the interaction phase of the experiment. Regions over each mouse’s whiskers and forelimbs are shown in magenta and cyan boxes, respectively, to estimate motion. B) Timeline of experimental paradigm (top) and ethograms for the stationary and moving mouse (bottom). C) Average percentage of time spent behaving during the first separated phase of the experiment (top) and the interaction phase (bottom). Intersecting regions show concurrent whisker and forelimb movements. D) Cross-correlation of each mouse’s binary behavior vectors during the interaction phase (black), compared with cross-correlation from the stationary mouse’s behavior vectors during the first and second separate phases (red). Behaviors across mice during the interaction phase were significantly correlated near 0 lag. E) Intersection over union for the behavior vectors was significantly greater during the interaction phase across mice than for the behavior vectors of the two separate phases from the stationary mouse. p<0.05; t-test.

### GCaMP Image Acquisition

GCaMP activity was imaged using Raspberry Pi Cameras (OmniVision OV5647 CMOS sensor) equipped with a triple-bandpass filter (Chroma 69013m). The lens on the camera has a focal length of 3.6mm with a field of view of ∼10.2×10.2mm, leading to a pixel size of ∼40 microns. 24-bit RGB images of GCaMP activity and reflectance were captured at ∼30fps and 256×256 resolution. The three cameras (2 brain and 1 behavior) were configured such that one camera was used to start the acquisition of the other two. Each cortex was illuminated using two LEDs simultaneously, where one light source (short blue, 447.5 nm Royal Blue Luxeon Rebel LED SP-01-V4 with Thorlabs FB 440-10 nm band pass filter) provides information about hemodynamic changes during the experiment (Xiao et al., 2017), and the other light source (long blue, 470nm Luxeon Rebel LED SP-01-B6 with Chroma 480/30 nm) excites GCaMP. The LEDs were turned on and off using a transistor-transistor logic (TTL) output from an isolated pulse stimulator (AM-Systems Model 2100) which was triggered immediately after the start of each experiment. This sudden change in illumination was used during post-hoc analysis to synchronize frames across cameras. With the current recording setup, the Raspberry Pi cameras occasionally drop frames as a result of writing the data to the disk. We identified the location of dropped frames by tagging each frame with a timestamp and found that consecutive frames were rarely dropped. Given the small number of dropped frames, and the relatively slow kinetics of GCaMP6s (T.-W. Chen et al., 2013), the lost data was estimated by interpolating the signal for each pixel, thus preserving the temporal resolution.

### GCaMP Image Processing

Image pre-processing was conducted with Python using a Jupyter Notebook (Kluyver et al., 2016). Further analysis was conducted using MATLAB (MathWorks, Natick MA, USA). Green and blue channels, which contain the GCaMP6s fluorescence and the blood volume reflectance signals respectively (Ma et al., 2016; Valley et al., 2020; Wekselblatt et al., 2016), were converted to ΔF/F_0_. The baseline image, estimated as the mean image across time for the entire recording, was subtracted from each individual frame (ΔF). The result of this difference was then divided by the mean image, yielding the fractional change in intensity for each pixel as a function of time (ΔF/F_0_).

To correct for hemodynamic artifacts, blue light (440+/-5nm) reflectance ΔF/F_0_ was subtracted from the green fluorescence ΔF/F_0_. In this way, small changes in the brain reflectance due to blood volume changes do not influence the epifluorescence signal. While we acknowledge that a green reflectance strobing and model-based correction may be advantageous (Ma et al., 2016), certain technical aspects of the Raspberry Pi camera (which is needed to perform this experiment due to its small form factor) such as its rolling shutter and inability to read its frame exposure clock prevent this method from being implemented. The short blue wavelength (440nm) is close to an oxy/deoxygenated hemoglobin isobestic point, and the reflected light signal correlates well with the green reflectance signal (Xiao et al., 2017), suggesting that this method effectively captures signal changes resulting from hemodynamic activity. Moreover, hemodynamic changes are relatively small compared to the signal-noise-ratio of GCaMP6s (Dana et al., 2014).

Occasionally, noisy extreme pixel values for ΔF/F0 were observed due to imaging near the edge of the window or due to the ratio-metric calculation of the ΔF/F0. To reduce their contribution, pixels exceeding a threshold value were set to be equal to the threshold, thereby reducing artifacts from smoothing or filtering that might result from inclusion of aberrantly large ΔF/F0 values. The threshold was set at the mean +/- 3.5x the standard deviation of each pixel’s time-series for GCaMP data, and at 15% ΔF/F0 for the reflectance data (which is larger than expected reflectance signal values). The ΔF/F0 signal was then smoothed with a Gaussian image filter (sigma=1) and filtered using a 4th order Butterworth bandpass filter (0.01-12.0Hz) (Vanni & Murphy, 2014).

### Behavior Quantification

To extract behavior events, a region of interest (ROI) was manually drawn on the behavior video over each mouse’s whiskers and forelimbs (Figure 2a). The motion energy within each ROI was calculated by taking the absolute value of the temporal gradient of the mean pixel value within the ROI. The resulting motion energy was smoothed via convolution with a Gaussian kernel (σ=5 frames) and a threshold was established at the mean + 1 standard deviation to detect behaviors. This analysis captured whisker and forelimb movements for the stationary mouse for the entirety of the experiment, and for the moving mouse only during the interaction phase (Figure 2c).

### Inter-brain correlation analysis

Correlation across brains was calculated using the Pearson’s correlation coefficient (PCC). To compare correlations across trial phases, the inter-brain PCC was calculated for a one-minute period during initial-separate, together, and final-separate trial phases. Global signals were calculated as the median ΔF/F_0_ across the entire dorsal cortex, whereas individual regions were selected from coordinates with respect to bregma according to the Allen Institute brain atlas, and the corresponding time-series data was calculated as the mean activity within a 5×5 pixel area surrounding each region location. Time-varying coherence between global signals was estimated with multitaper methods over a 45 second window with 22.5 second overlap using the Matlab Chronux toolbox with a time-bandwidth product of 5 and a taper number of 9 (Bokil, Andrews, Kulkarni, Mehta, & Mitra, 2010; Mitra & Bokil, 2009).

### Whisker triggered event analysis

Calcium activity surrounding whisk-initiation events (+/-1s) were extracted from each whisk event and averaged across events per trial. Whisk events were excluded if any of the following conditions were met: 1) they occurred coincidentally with forelimb movements, 2) they occurred within 1 second of the previous whisk event, or 3) total duration exceeded 0.5 seconds. Self-initiated whisking activity therefore refers to averaged calcium activity of mouse A surrounding whisk events initiated by mouse A, whereas partner-initiated maps refer to averaged calcium activity of mouse A surrounding whisk events initiated by mouse B.

### Statistics

Statistical tests were conducted with MATLAB. All data were tested for normality using a Kolmogorov-Smirnov test prior to subsequent statistical analyses. Correlation values were transformed using Fisher’s z-transformation. Comparisons between two groups were conducted using two-tailed t-tests for parametric data and Wilcoxon signed rank tests for non-parametric data. Comparisons between trial phases were assessed using a repeated measures ANOVA with post-hoc Bonferroni correction for multiple comparisons. All statistically significant results were observed on the GCaMP signals alone as well as the hemodynamic corrected signals.

### Resource Availability

Resources to assist in building cortex-wide GCaMP imaging systems, including parts lists, assembly instructions, and CAD files are available in the supplemental information and at the Open Science Framework project entitled Dual Brain Imaging. Code for image acquisition, preprocessing, and analysis are available at the University of British Columbia’s Dynamic Brain Circuits in Health and Disease research cluster’s GitHub (https://github.com/ubcbraincircuits/dual-mouse). Data are available in the Federated Research Data Repository at https://doi.org/10.20383/101.0303.

## Results

### Mice Exhibit Correlated Bouts of Behavior During Social Interaction

Forelimb and whisker movements were monitored for each mouse to measure behavior (Figure 2A) using a camera positioned underneath the interacting mice (Figure 1C). The stationary mouse’s behavior was captured throughout the entire experiment, while the moving mouse’s behavior acquisition was limited to the social interaction period only (Figure 2B).. Bouts of forelimb and whisker movements often occur simultaneously (Figure 2C), and the amount of time spent actively moving whiskers or forelimbs, expressed as percentage of time spent behaving in each trial phase, did not change between the separate period and the interaction period (Figure 2C, n=33 trials, 14.1+/-3.4% whisking separate vs 14.4+/-4.3% whisking interaction, and 8.9+/- 3.8% forelimb separate vs 9.8+/-4.8% forelimb interaction period, n=33 trials, p=0.77, paired t-test). The total number of behavior events did not differ across trial phases (Supplemental figure 1a-b). Behavior periods across mice exhibited temporal coordination, as shown by the peak in the cross-correlation of binary behavior vectors at time lag 0s (Figure 2D black). This temporal relationship was compared with the cross-correlation of the two behavior vectors measured from the stationary mouse during the separate phases of the experiment (Figure 2D, red). The correlation at 0 lag was significantly greater for the inter-animal behaviors compared to the two separate epochs from the stationary mouse (Figure 2d, n=33 trials, 0.32 +/- 0.11 together vs 0.17+/- 0.11 separate, p=6.1×10^−8^, paired t-test), as well as any other combination of epochs that included at least one separated period (Supplemental figure 1c). Intersection over union for the two behavior vectors (a measure of shared behavior) was significantly greater across animals than across the separate epochs (Figure 2e, n=33 trials, 0.19 +/- 0.09 together vs 0.09+/-0.07 separate, p=1×10^−6^, paired t-test).

### Global Calcium Signals Synchronize During Interaction

Global signals were calculated as the spatially averaged ΔF/F_0_ across the entire masked region of the two cortical hemispheres (Figure 3A-B). Global signal synchronization, measured as the Pearson’s correlation coefficient (PCC) between global signals from each mouse, was significantly higher during the interaction phase (0.19 +/- 0.23 interaction phase) of the experiment than during either of the two separate phases (−0.007 +/- 0.13 before; −0.01 +/- 0.14 after). (Figure 3C, repeated measures ANOVA, n=35 trials, trial-phase: F _2,68_ = 11.2, p = 0.002=). Cagemate vs non-cagemate pairings did not have a significant effect on inter-brain correlation (repeated measures ANOVA, F_1,34_ = 6×10^−4^, p = 0.99). This increase in inter-brain correlation was not observed in experiments using the Thy1-GFP mouse line (Feng et al., 2000), suggesting that hemodynamic contributions to the fluorescence signal do not account for this synchronization (Supplemental Figure 2). Additionally, inter-brain PCCs during the interaction phase were significantly higher than PCCs observed across trial-shuffled global signal pairings during the interaction phase (Figure 3C, 0.02 +/- 0.14, n=35 correct pairs vs n=595 shuffled pairs, p=1.2×10^−11^, t-test), suggesting that inter-brain synchronization is interaction-specific and cannot be attributed to environmental variables shared across trials e.g. timing of the stage translation. An expanded view of the interaction period from Figure 3B is shown with behavior annotations in Figure 3D. Prominent calcium events are often accompanied by sustained periods of behavior (Figure 3D) and the average ΔF/F_0_ during the behavior event was positively correlated with behavior duration (Figure 3E, n=2075 behavior events, r=0.44, p<0.001). The time-varying coherence between brains revealed an increase in global signal coherence during the interaction phase at frequencies below 0.2Hz (Figure 3F), suggesting that the large, low-frequency calcium signals observed during behavior periods drive the interbrain global signal synchronization.

**Figure 3.**
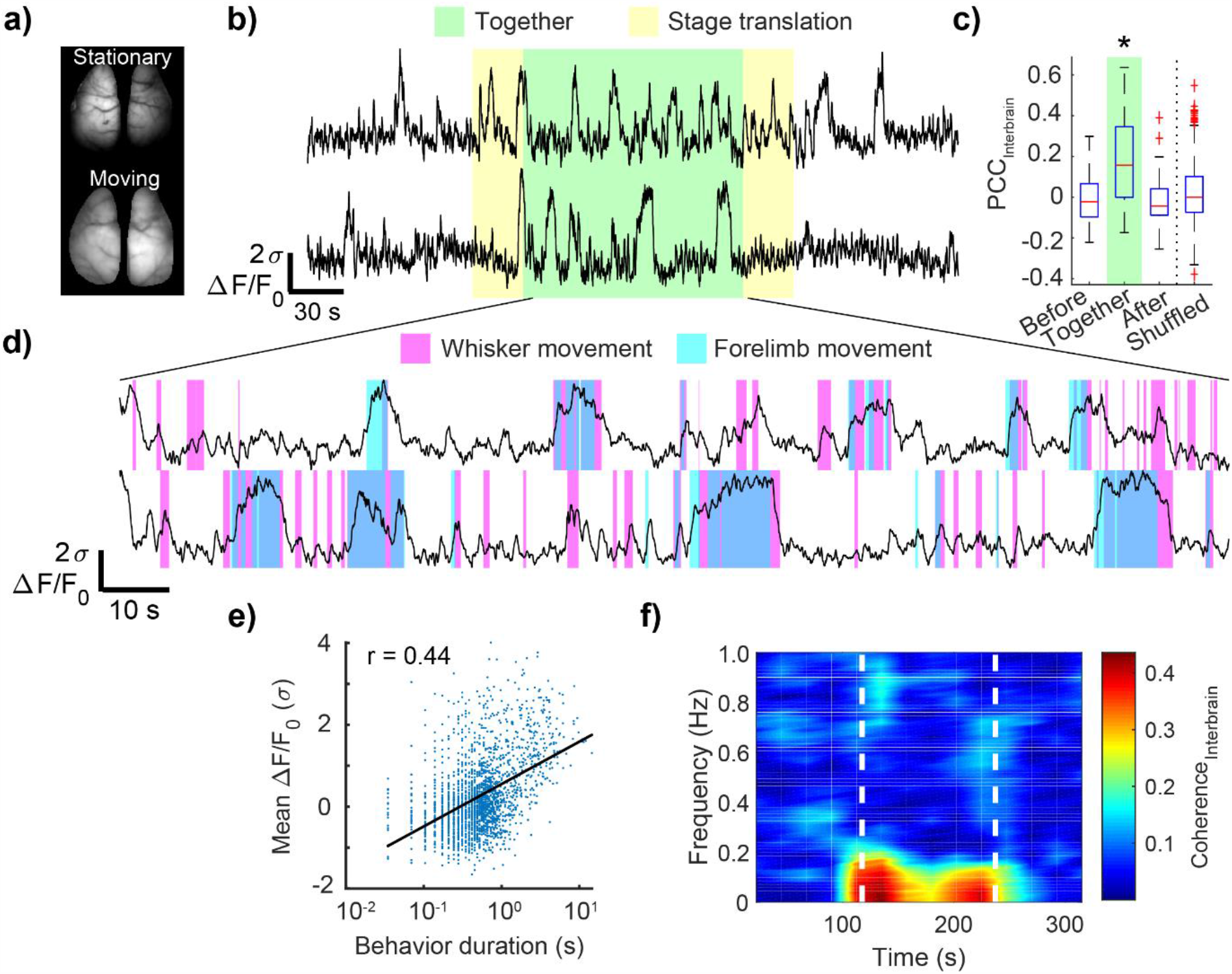
Global signal synchronization during mouse interaction. A) Example images of dorsal cortical windows for each mouse. B) Representative example of GCaMP6s activity averaged across the cortical mask for each mouse. C) Pearson correlation coefficients of the two global signals were significantly greater during the interaction phase than either of the two separate phases (p<0.001; repeated measures ANOVA with post-hoc Bonferroni correction for multiple comparisons). Inter-brain correlations during interaction were significantly greater than trial-shuffled interaction-phase pairings (p<0.001; t-test). D) Expanded view of global signals during interaction phase with behavior annotations overlaid. E) Global signal ΔF/F_0_ is positively correlated with duration of behavior (Pearson correlation coefficient; r=0.44; p<0.001). F) Time-varying inter-brain coherence, computed with a 45s window and averaged across all experiments, shows an increase in coherence from 0-0.2Hz during the interaction phase (white dashed lines).

### Inter-brain synchronization across multiple cortical regions during social interaction

To determine whether the increase in cortex-wide inter-brain synchronization during social interaction was a global phenomenon or instead attributed to specific cortical regions, we examined inter-brain correlations on a region-by-region basis, including motor areas (vibrissae motor cortex, vM1; secondary motor cortex, M2; anterior lateral motor cortex, ALM), sensory areas (primary visual cortex, V1; forelimb, FL; hindlimb, HL, and anterior and posterior barrel cortex, aBC and pBC), retrosplenial, (RS), and parietal association area (PTA) (Figure 4A). All selected regions showed a relative increase in inter-brain correlation during the interaction period (Figure 4B-C, 0.17+/-0.03 together vs -0.013+/-0.009 before and -0.019+/-0.02 after interaction; repeated measures ANOVA, n=35 trials, F_2,18_ = 53.8, p = 8.1×10^-5^,), with the most dramatic increase observed from pBC. Intra-brain correlations also showed a slight but significant increase during the interaction period (Supplemental Figure 3, n=35 trials, p<0.05, repeated measures ANOVA with post-hoc Bonferroni correction).

**Figure 4.**
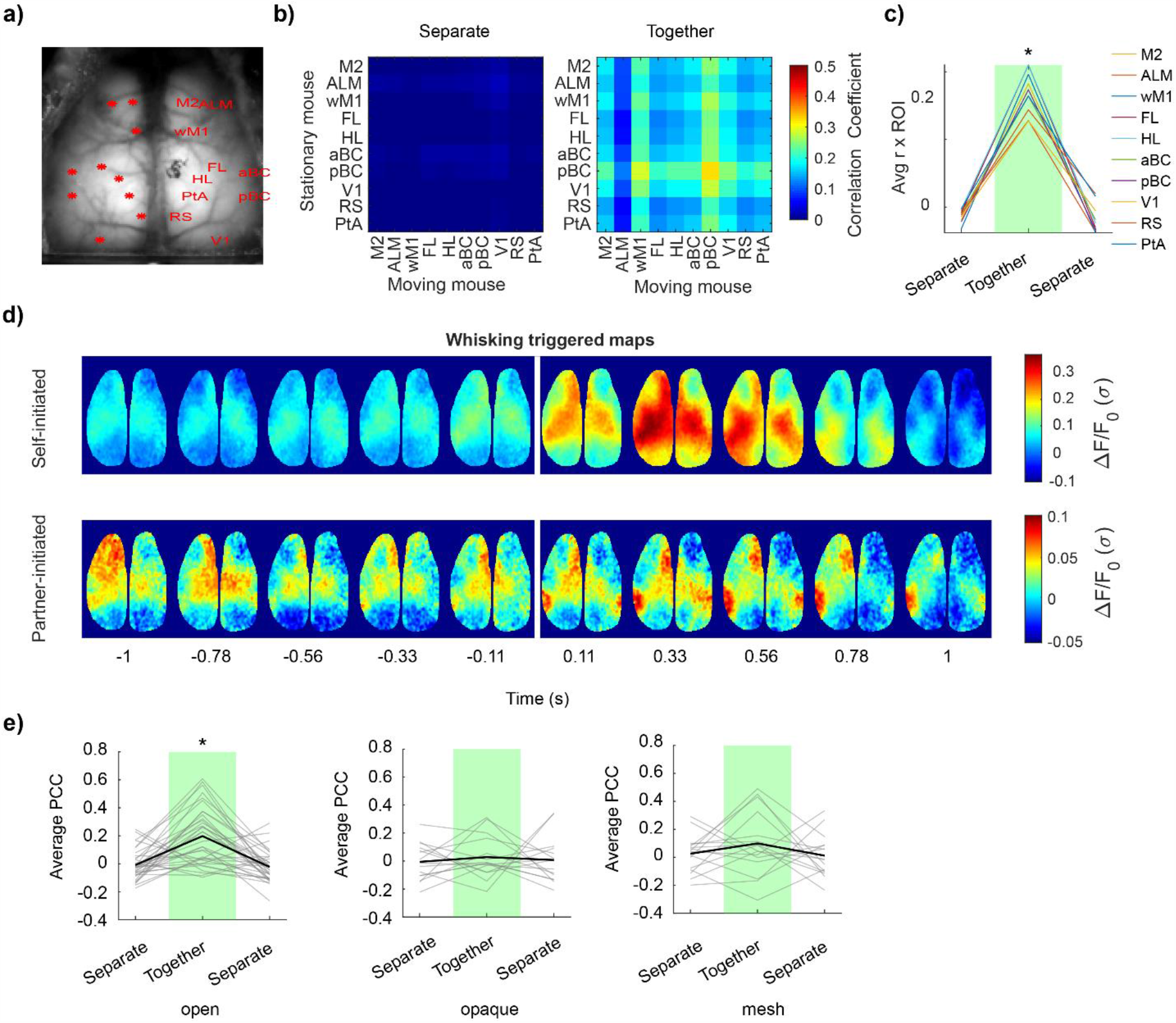
Whisker contact drives multi-region inter-brain synchronization. A) Example image of transcranial mask with regions labelled. Abbreviations: ALM, anterior lateral motor cortex; M2, secondary motor cortex; vM1, vibrissae motor cortex; aBC, anterior barrel cortex; pBC, posterior barrel cortex; HL, hindlimb; FL; forelimb; lPTA; lateral parietal association area; RS, retrosplenial cortex; V1, primary visual cortex. B) Averaged inter-brain correlation matrices across all experiments during the period before interaction (left) and the period during interaction (right).C) Average inter-brain correlation for each region of interest against all other regions, averaged across mice (*p<0.05; repeated measures ANOVA with post-hoc Bonferroni correction). Error bars not shown for clarity. D) Example montage showing whisker movement triggered activity. E) Averaged region-by-region correlations in each trial phase for open interaction experiments (left, n=35) and barrier controls (middle, n=15; right, n=16). Individual trials are represented by thin grey lines, and trial averages are represented by thick black lines (*p<0.001, repeated measures ANOVA with post-hoc Bonferroni correction).

Given the relatively large increase in inter-brain correlation observed from the barrel cortex areas, we wondered if shared sensory experiences during social tactile investigation could underlie the increase in inter-brain correlation. We examined self-initiated and partner-initiated whisking montages, which show cortical dynamics surrounding whisking bout initiation events. Averaged self-initiated and partner-initiated montages showed activation of the posterior barrel cortex area (Figure 4D), confirming that whisking activity elicited by either mouse can elicit barrel cortex responses in both mice (i.e. a shared sensory experience). To test if this shared sensory experience was important for establishing the inter-brain correlation, we performed experiments with physical barriers in place to prevent whisker-whisker contact between mice. Significant increases in region by region inter-brain correlation were not observed when physical contact was prevented using an opaque cardboard sheet or a transparent copper mesh (Figure 4e, open trials: 0.17+/-0.19 together vs −0.007+/-0.11 before and −0.005+/-0.11 after interaction; repeated measures ANOVA, n=35 trials, F_2,68_ = 13.95, p=0.0007; opaque trials: −0.023+/-1.3 together vs − 0.024+/-0.09 before and 0.03+/-1.3 after, repeated measures ANOVA, n=15, F_2,28_ = 1.0, p = 0.34; mesh trials: 0.09+/-0.22 together vs −0.011+/-0.11 before and −0.012+/-0.11 after, repeated measures ANOVA n=16, F_2,30_ = 0.022, p = 0.88). Cagemate vs non-cagemate effects were not significant (open trials: F_1,34_ = 3.2×10^−5^, p = 1.0; opaque trials: F_1,13_ = 0.62, p = 0.45; mesh trials: F_1,14_ = 0.63, p = 0.44). Furthermore, no ultrasonic vocalizations were observed during these experiments (Supplemental Figure 4).

### Cortical activation during whisking on familiar partner is stronger than non-familiar partner

To determine if partner identity has an effect on cortical activity, we examined the relationship between social rank and social novelty on cortical dynamics. Social rank differences between cagemate partners, as determined by the tube test assay, were not strongly correlated to the magnitude of global signal inter-brain synchrony at the cortex-wide scale (Figure 5A, n=14 trials, Pearson’s r=0.15, p=0.6). Similarly, no differences in inter-brain synchronization were seen when comparing interactions between cagemates vs interactions between non-cagemates (Figure 5B, cagemates n=16 trials, 0.18 +/- 0.23; non-cagemates, n=7 trials, 0.21 +/- 0.24; t-test; p=0.66). However, self-initiated whisking events produced greater global cortical activation during cagemate trials compared to non-cagemate trials (Figure 5C-E, cagemates n=40 trials, 0.09+/-0.19; non-cagemates, n=26 trials, −0.07+/-0.23; p=0.002, t-test). This difference could not be attributed to differences in the number or duration of whisking bouts (Figure F-G, number of whisking events: 12.1 +/- 4.3 cagemates vs 11.3 +/- 3.4 non-cagemates, p=0.4, t-test; duration of whisking events, 0.27 +/- 0.08 s cagemates vs 0.27 +/- 0.06, p=0.1.0, t-test).

**Figure 5.**
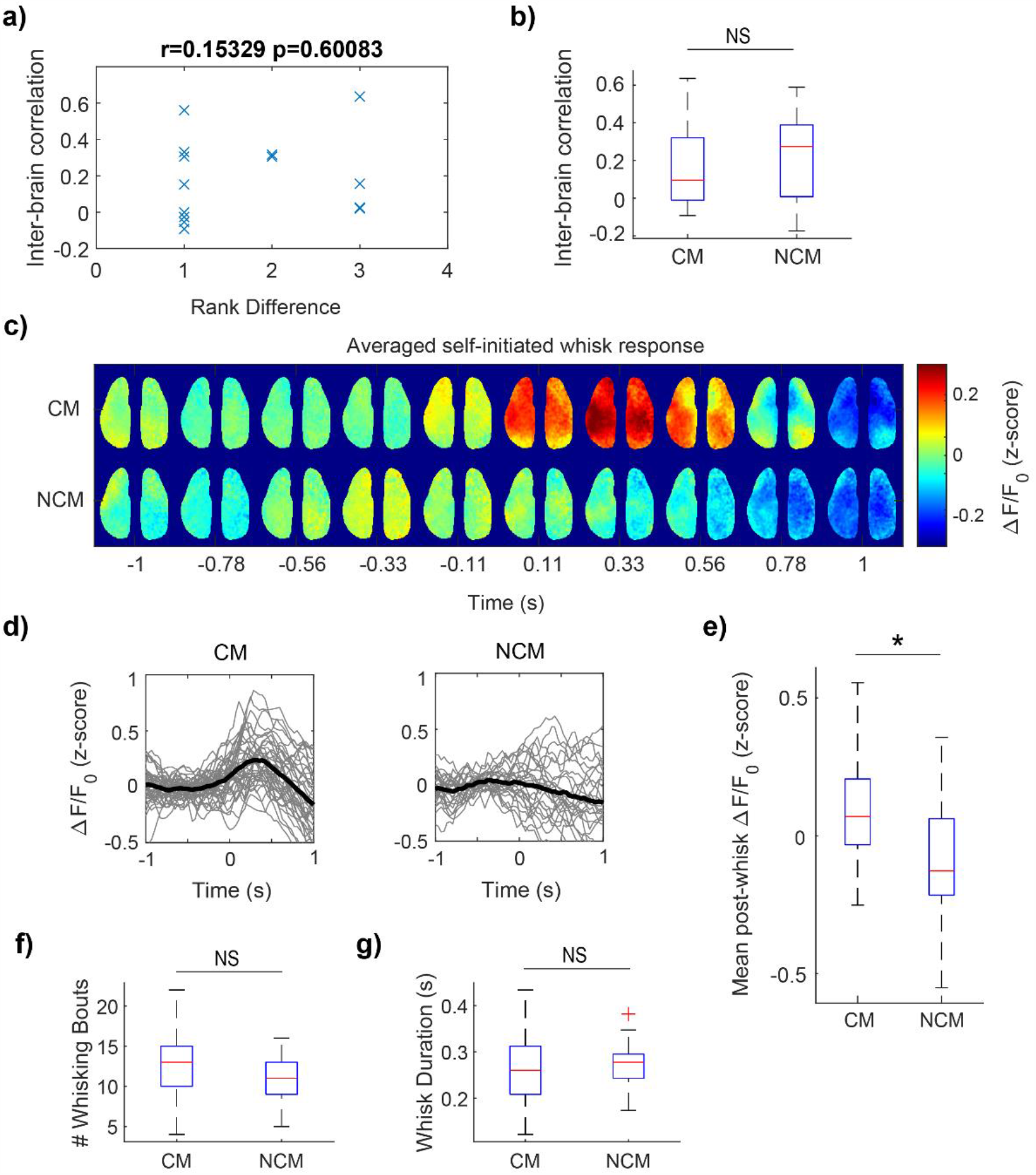
Widespread cortical activation during whisking on a familiar versus novel partner. A) Inter-brain global signal correlation is not related to social rank differences between interacting partners (Pearson correlation coefficient, r=0.15). B) Inter-brain signal correlation does not depend on cagemate vs non-cagemate experiments (t-test, p=0.66). C) Self-initiated whisk event triggered activation maps for cagemates and non-cagemates, averaged across all whisk events per mouse then averaged across mice. D) Global signals associated with whisk-triggered events for cagemates and non-cagemates. Thin gray lines show averaged global signals across all whisking events per mouse. Thick black lines show average across mice. E) The mean ΔF/F_0_ in the 1-s post-whisk period was significantly greater in the cagemate group compared to the non-cagemate group (t-test, p=0.005). F-G) Number of whisking bouts and duration of whisking bouts were not different between cagemate and non-cagemate groups.

## Discussion

Our results indicate widespread correlated cortical activity between the brains of interacting mice. This synchrony is not associated with the mechanics or timing of the imaging paradigm as it was not present when trial-shuffled mouse pairs were examined. Rather, the inter-animal cortical synchronization is likely driven by temporally coordinated bouts of behavior (e.g. whisking or forelimb movements) and shared somatosensory experiences. Although previous work found that the magnitude of inter-brain synchronization may convey information regarding the social status of one of the individuals (Jiang et al., 2015; Kingsbury et al., 2019), we did not find a relationship between social rank differences and cortex-wide inter-brain synchronization. Surprisingly, we found that whisking behavior in the presence of a familiar conspecific partner elicited more pronounced cortical activation compared to whisking onto a non-familiar partner. This difference may suggest a macroscale cortical network representation of social partner identity. Future work will examine the contribution of social sensory cues on cortex-wide behavior and individual sensory circuits.

One limitation of the presented work is that the frame rate of the behavior camera was not fast enough to clearly resolve whisker movements. Detailed analyses of whisker movements in mice typically use camera acquisition rates of ∼500fps (Mayrhofer et al., 2019; Sofroniew, Cohen, Lee, & Svoboda, 2014). It is possible that some whisking events were missed by our analyses, or the precise timing of whisk initiation was not accurately resolved. Nevertheless, whisker motion energy measurements resolved the initiation of gross whisking events, as suggested by the cortical maps displaying barrel and vibrissae motor cortex activation (Figure 4d); and false-negative error rates should presumably be consistent across experiments. Another limitation of the presented work is that mice must be head-restrained in order to be imaged and positioned properly. In a previous study, head-fixation was found to be aversive, but with training and habituation stress recedes (Z. V Guo et al., 2014) and rodents can even be trained to restrain themselves (Aoki, Tsubota, Goya, & Benucci, 2017; Murphy et al., 2020; Scott, Brody, & Tank, 2013). For this reason, we present the results as an interaction that occurs in the context of head-fixation and caution that the observed brain dynamics may not reflect true naturalistic social touch behavior. Despite this, head-restraint facilitates consistent and reproducible interactions between animals, allowing for trial-averaging of behaviors. Recent development of a head-mounted mesoscopic camera allows for the exciting possibility to examine cortex-wide neural dynamics during more naturalistic social interactions in freely-moving mice (Rynes et al., 2020).

In conclusion, we introduce a dual mouse mesoscale imaging platform that can create reproducible interactions between mice that constrains some of the possible behaviors and timing due to the head-restrained and rail-based system. Such a constraint may be particularly important when evaluating the behavior of different mouse mutants associated with disorder of social interactions, such as the SHANK3 mutant mice (Peça et al., 2011). Future experiments can incorporate simultaneous electrophysiological recordings (Xiao et al., 2017) or examine lower frequency events that are revealed using functional near-infrared spectroscopy or intrinsic optical signals to draw parallels to human studies analyzing inter-subject interactions.

## Acknowledgements

We thank Pumin Wang for help with surgery and Matthieu P. Vanni, Allen W. Chan, Dongsheng Xiao and Alexander McGirr for helpful discussion and comments.

