## Supplemental figures for "Dual brain cortical calcium imaging reveals social interaction-specific correlated activity in mice"

**Supplemental Video 1. Sliding window correlation and global signal response for interacting mice.** Data from two head-restrained tTA-GCaMP6s mice at the onset of the interaction phase. Time is shown in frames. The two pseudo-colored images show the processed calcium images from the left mouse (top) and right mouse (bottom) over the 8mm field of view of the dorsal cortex. The colorbar indicates the $\Delta F/F_0$ of the signals.

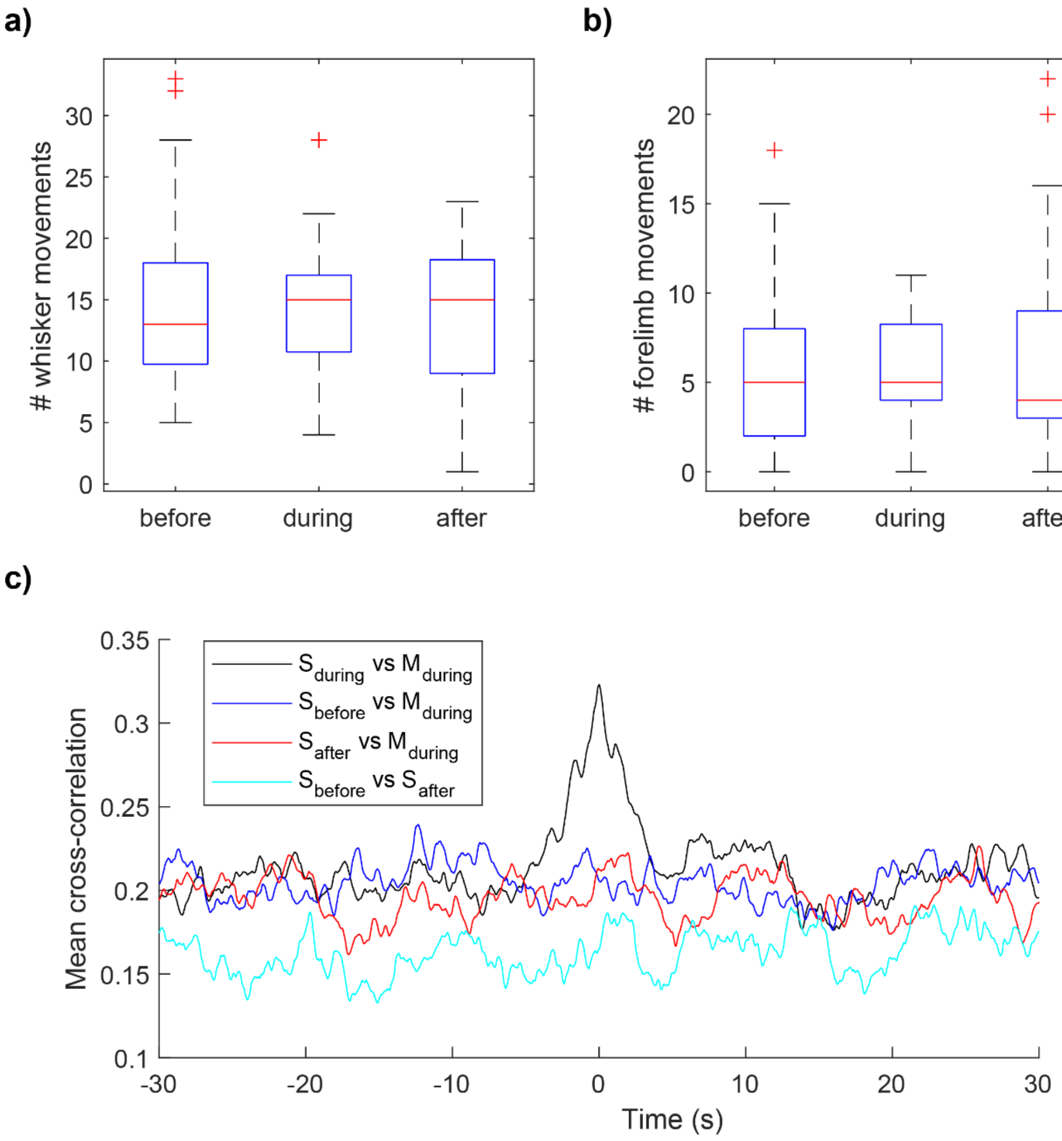


**Supplemental Figure 1: Whisker and forelimb movements across trial phases.** A-B) number of whisker movements (a) and forelimb movements (b) from the stationary mouse across all trial phases (one minute period). C) Cross-correlation results for behavior vectors (forelimb and whisker combined) mismatched across trial phases. No discernable peaks are seen in any mismatched comparison. Legend abbreviations: S – stationary mouse, M – moving mouse, during – interaction period, before/after – before/after interaction period.

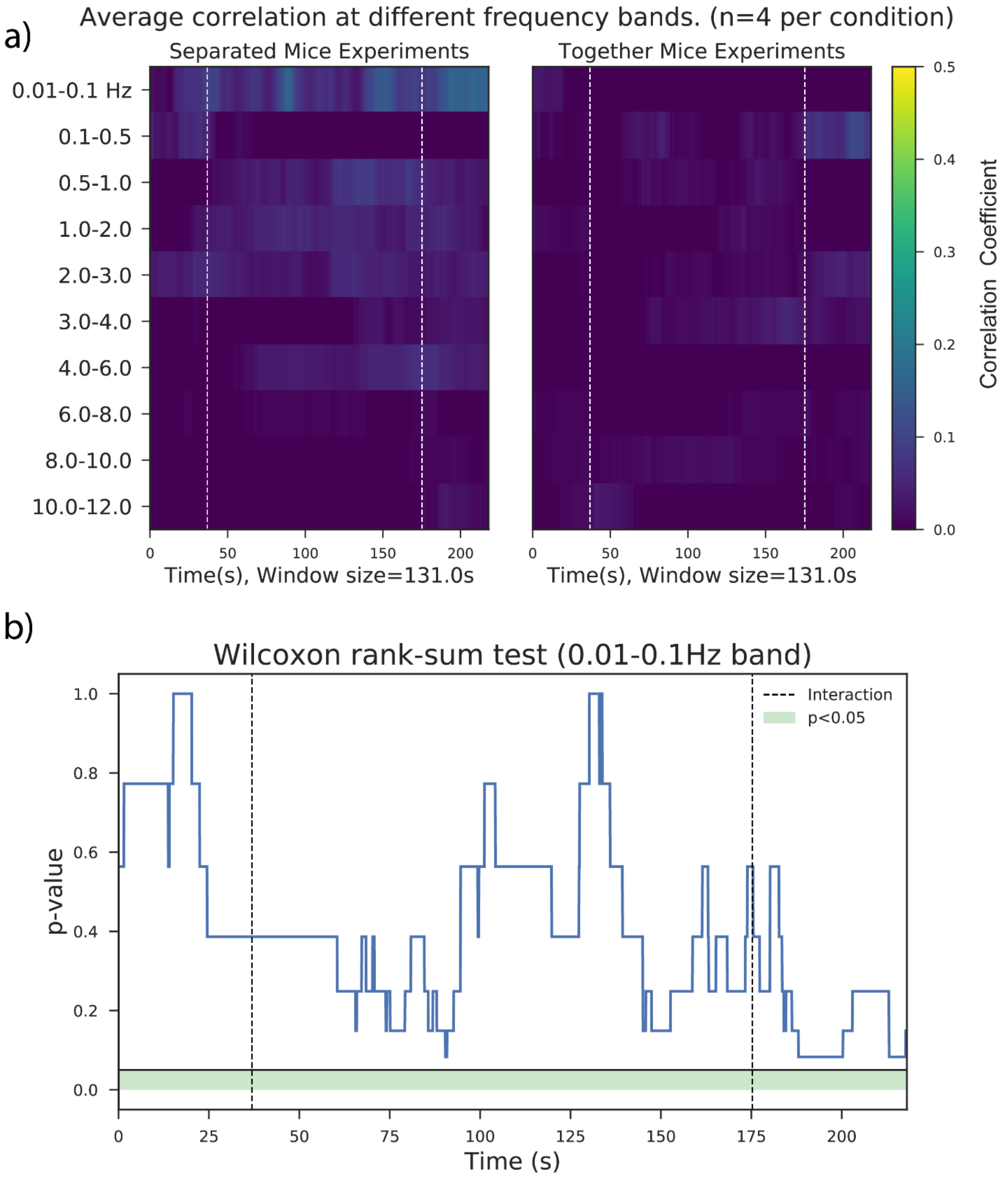


**Supplemental Figure 2. Thy1-GFP Mice do not Exhibit Global Signal Synchronization.** A) Global signals from Thy1-GFP mice were filtered at 10 different frequency bands, and the Pearson correlation coefficient vs time was computed using a sliding window correlation. B) P-values for sliding window correlation between trials where mice were fully separated for the entire experiment vs trials where mice were brought together.

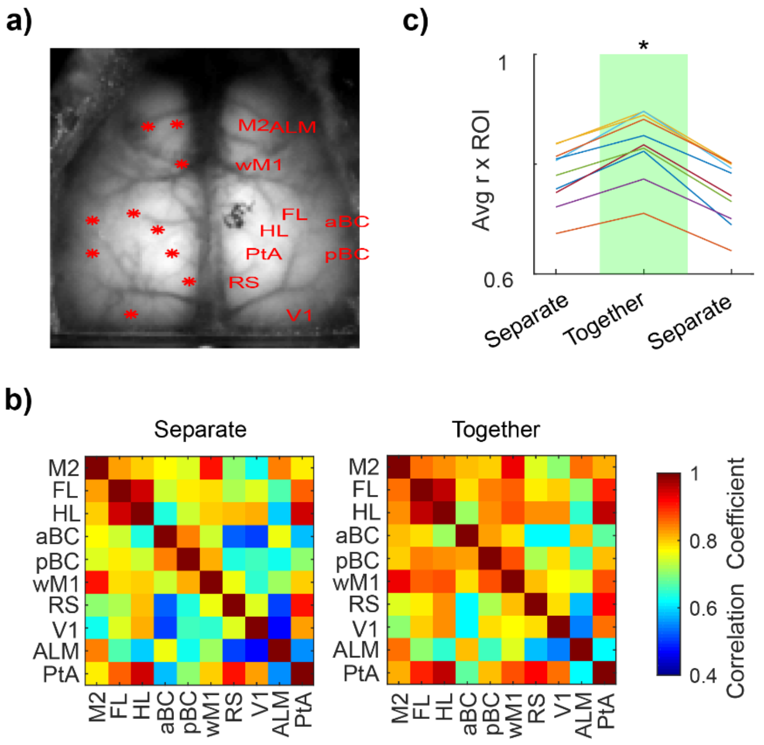


**Supplemental Figure 3. Correlation maps intra-brain group data.** A) Cortical fluorescence image with analyzed regions of interest overlaid. B) Within brain correlations matrices for the specified regions. C) Intra-brain correlation increases slightly during the together phase.

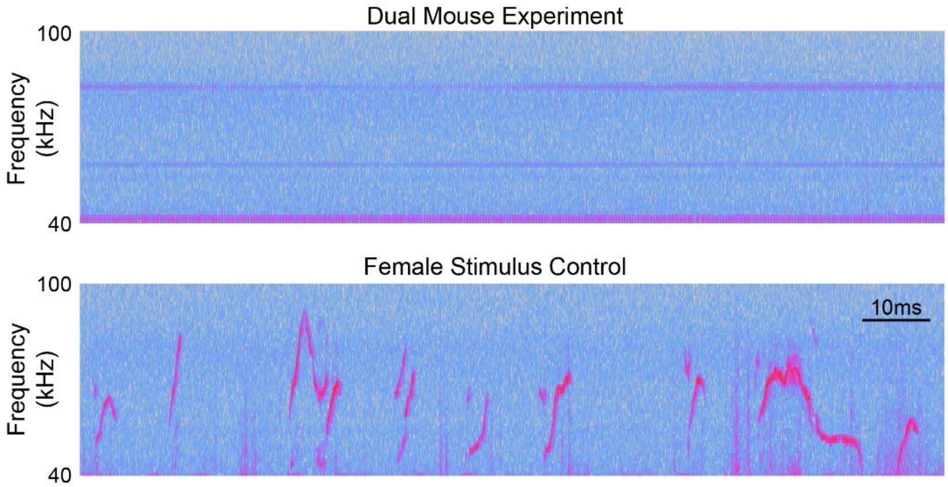


**Supplemental Figure 4. No ultrasonic vocalizations detected in social-interaction tests.** Example data from the social interaction experiment (top), compared to a control experiment taken from a breeder mouse introduced to a female (bottom). Ultrasonic vocalizations are clearly observed in the female stimulus control experiment, but not in the two-mouse imaging experiments.
